# Automated microscopic measurement of fibrinaloid microclots and their degradation by nattokinase, the main natto protease

**DOI:** 10.1101/2024.04.06.588397

**Authors:** Justine M. Grixti, Chrispian W. Theron, J. Enrique Salcedo-Sora, Etheresia Pretorius, Douglas B. Kell

## Abstract

Nattokinase, from the Japanese fermented food natto, is a protease with fibrinolytic activity that can thus degrade conventional blood clots. In some cases, however, including in Long COVID, fibrinogen can polymerise into an anomalous amyloid form to create clots that are resistant to normal fibrinolysis and that we refer to as fibrinaloid microclots. These can be detected with the fluorogenic stain thioflavin T. We describe an automated microscopic technique for the quantification of fibrinaloid microclot formation, which also allows the kinetics of their formation and aggregation to be recorded. We also here show that recombinant nattokinase is effective at degrading the fibrinaloid microclots *in vitro*. This adds to the otherwise largely anecdotal evidence, that we review, that nattokinase might be anticipated to have value as part of therapeutic treatments for individuals with Long COVID and related disorders that involve fibrinaloid microclots.

## 1. Introduction

Thrombosis, the blocking of blood vessels by blood clots, along with the related thrombo-inflammation and thromboembolism, is a chief cause of cardiovascular disease [1-6]. Consequently anything that can promote safe anti-coagulation or fibrinolytic activity is likely to have therapeutic potential (e.g. [7-15]).

Nattō (usually rendered natto) is a Japanese food made via the fermentation of soy beans using the Gram-positive organism *Bacillus subtilis* var natto [16-20]. It has been widely consumed for over 2000 years, and is considered safe [21]. The proteolytic activity of natto was detected in 1906 [22] and its fibrinolytic activity in 1925 [23]. However, it was not until 1987 [21] that an enzyme exhibiting these activities was purified from natto; in spite of it being a protease it was termed nattokinase [21].

Despite having to pass through the gut wall [24-31], nattokinase is orally available (and this can be improved [32-34]), is considered a major contributor to the purported health benefits of natto [21,27,35-60], and is itself recognised as safe [61-63].

The experimental 3D structure of nattokinase, which is a serine protease related to subtilisin, is available [64,65], and it may also be produced via purification [66-69] or (as here) recombinantly [44,70-83]. Although not our prime focus in this paper, it is also known to cleave plasminogen activator inhibitor I [84], to have antiplatelet [85], anti-inflammatory [86], and anti-hypertensive [87-89] activities, and to show neuroprotective [90] and post-stroke benefits [91] as well, when dosed adequately, as having anti-lipidaemic effects [92].

Following earlier work using electron microscopy (e.g. [93-96]), we discovered that fibrinogen could polymerise or clot into an anomalous, amyloid form of fibrin (e.g. [97-104]) that exactly reflected the clots seen in both the electron microscope [105] and in bright field optical microscopy [106]. As with prions and other amyloid forms of proteins [98,107], that are often highly resistant to proteolysis (e.g. [108,109]), the existence of these ‘fibrinaloid’ microclots implies their comparative resistance to normal fibrinolysis [110,111], with their precise structures [112] being affected by other small and macromolecules and ions that they may have bound [97,103,113-119]. The varieties of stable macrostates into which a given amyloidogenic sequence can fold (even under the same conditions [120,121]) are referred to as different ‘strains’ [122-132] or ‘polymorphisms’ [133-144], and in some cases are sufficiently stable (i.e. kinetically isolated from other macrostates) that they are even heritable [122,145-151]. Homo- and hetero-polymerisation and their catalysis are then referred to, respectively, as (self-)’seeding’ [140,152-166] and ‘cross-seeding’ [153,167-174]. More recently, we have established the prevalence of these fibrinaloid microclots in post-viral diseases such as Long COVID [106,175-178] (and see [179]) and ME/CFS (myalgic encephalopathy/chronic fatigue syndrome) [180,181]. The lower amyloidogenicity of omicron versus earlier variants of SARS-CoV-2 is also reflected in its lower virulence [182], implying that these microclots are on the aetiological pathway of the disease, and they can explain many symptoms [183], including fatigue [184], post-exertional symptom exacerbation [185], autoantibody generation [107] and Postural Orthostatic Tachycardia Syndrome (POTS) [186]. Fibrin amyloid microclots also occur during sepsis [187], while amyloid deposits are also observed in the skeletal muscles of those with Long COVID [188]. Overall, this ability of fibrinaloid microclots to provide a mechanistic explanation of multiple phenomena is consistent with the ‘explanatory coherence’ view of science [189-192]. In common with other amyloid proteins [98], that contain a characteristic cross-β motif [173,193-209], they can be visualized using the fluorogenic stain thioflavin T [144,173,210-224] or via vibrational spectroscopy [221,225-236]. As with any other ligand or binding agent, the rotation of the bound form is more restricted than that of the free form (which is largely what makes it fluorogenic), and precise intensities of thioflavin T fluorescence depend on the location and conformation(s) to which the thioflavin T is bound [212-214,237-254] and in some cases on the presence of interferents [255].

Although nattokinase preparations are widely available commercially, and as noted above they are considered to have significant therapeutic value, including in Long COVID [256,257], their exact contents are uncertain, and so we decided that it was best to create and use purified, recombinant material.

While the proteolytic specificity of nattokinase, as an alkaline serine protease [44,258,259], is surprisingly underexplored, beyond a broad similarity to that of plasmin [44,260] (and nattokinase can even degrade spike protein [261] and certain ‘classical’ amyloids [262-264]), the question arises as to whether or not nattokinase can degrade <u>the amyloid ‘fibrinaloid’</u> form of microclots. The purposes of this paper are (i) to describe an efficient, quantitative, automated microscopic method that can be used to determine the size and number of amyloid microclots and any time-dependent changes therein, and thus (ii) to assess any such nattokinase-induced degradation of the microclots, concluding that nattokinase can indeed degrade fibrinaloid microclots effectively. The therapeutic implications of this are discussed.

## 2. Results

### Basic phenomenon, and effect of concentration of NK and incubation time

To give an indication of the kinds of data obtained in this study, Figure 1 (left panels) shows three Cytation images representing clots as stained with thioflavin T following incubation of fibrinogen plus thrombin plus LPS (as in [97]) for 6h, either with no further additions (Figure 1A, top), with PBS (Figure 1B, middle), or after simultaneous exposure to 28 ng/mL nattokinase in PBS (Figure 1C, bottom). In addition, the right hand panels of the figure show the 8-bit intensity distributions of pixels on linear and logarithmic scales.

**Figure 1.**
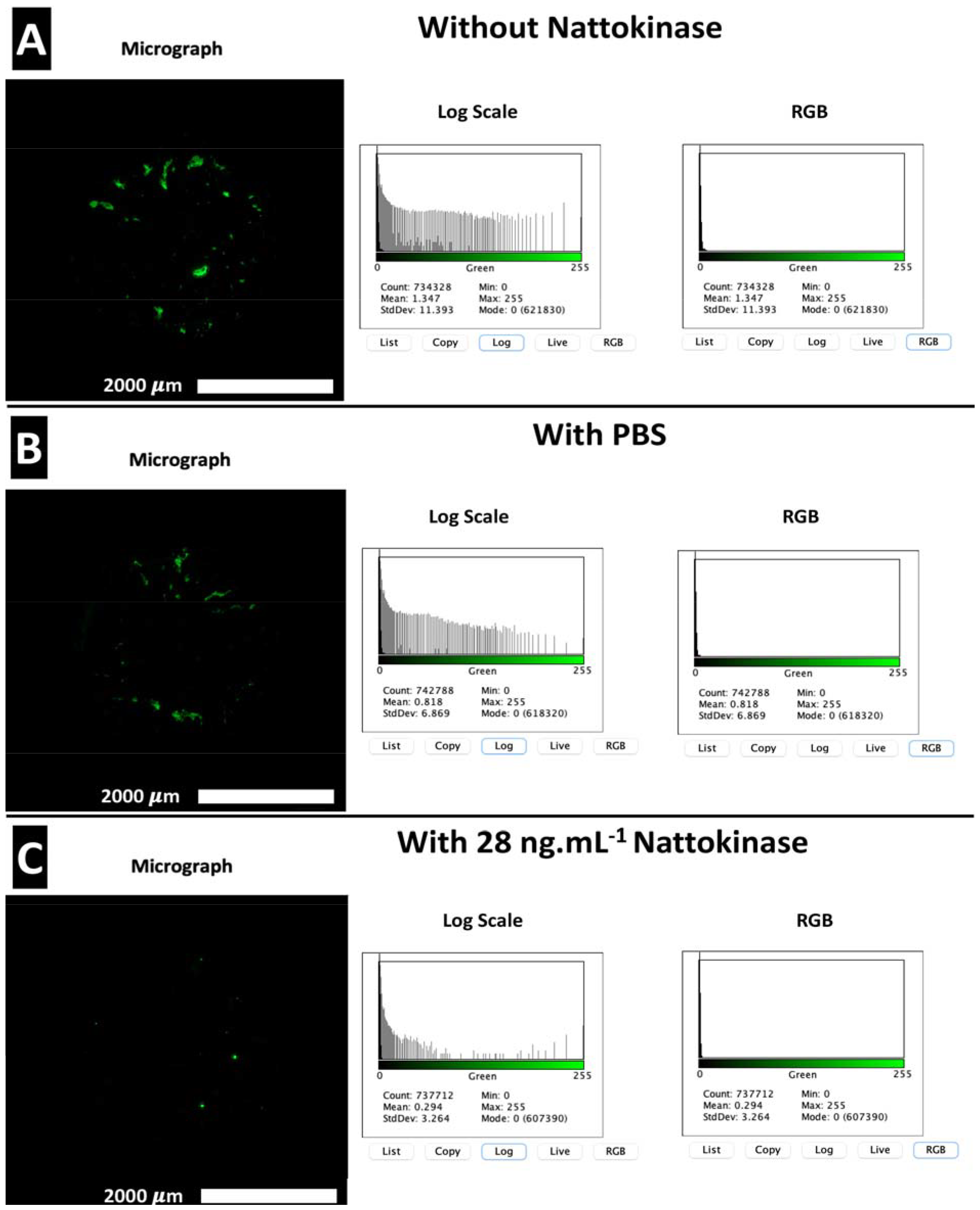
Images of fibrinaloid microclot formation and their removal via nattokinase. Thrombin and fibrinogen were incubated together with thioflavin T and LPS, and imaged after 6h in a Cytation 1, as described in Methods. Further additions were (A) none, (B) PBS, (C) recombinant nattokinase 28 ng/mL. Bar = 2 mm (2,000 μm).

While we sought to avoid any ‘cherry picking’ in the past, the great attraction of the present approach is that the entire sample is imaged (serially) so this issue is completely avoided. Although not necessarily obvious to the naked eye, there are variations in pixel intensity that allow a thresholding to determine what counts as a clot boundary. Figure 1 also shows the pixel intensity variation for the images displayed on its left side; the logarithmic plot in particular makes clear how much the pixels of larger intensity differed following the addition of the nattokinase.

The time evolution of these data (Figure 2) shows that in the absence of nattokinase the clot numbers increase for an hour or so then decrease slightly before stabilizing (Figure 2A). When nattokinase is present the clot numbers decrease after the first time point and by 2h have attained their lowest level, this being approximately half that of the 14 ng/mL nattokinase (in which the nattokinase level is thus halved), possibly implying a loss of activity over time. In Figure 2B we see the dynamics of the clot intensity (total number of pixels), this being substantially lower in the presence of NK, especially at the higher level of enzyme. Figure 2C shows the time evolution of the median clot size.

**Figure 2.**
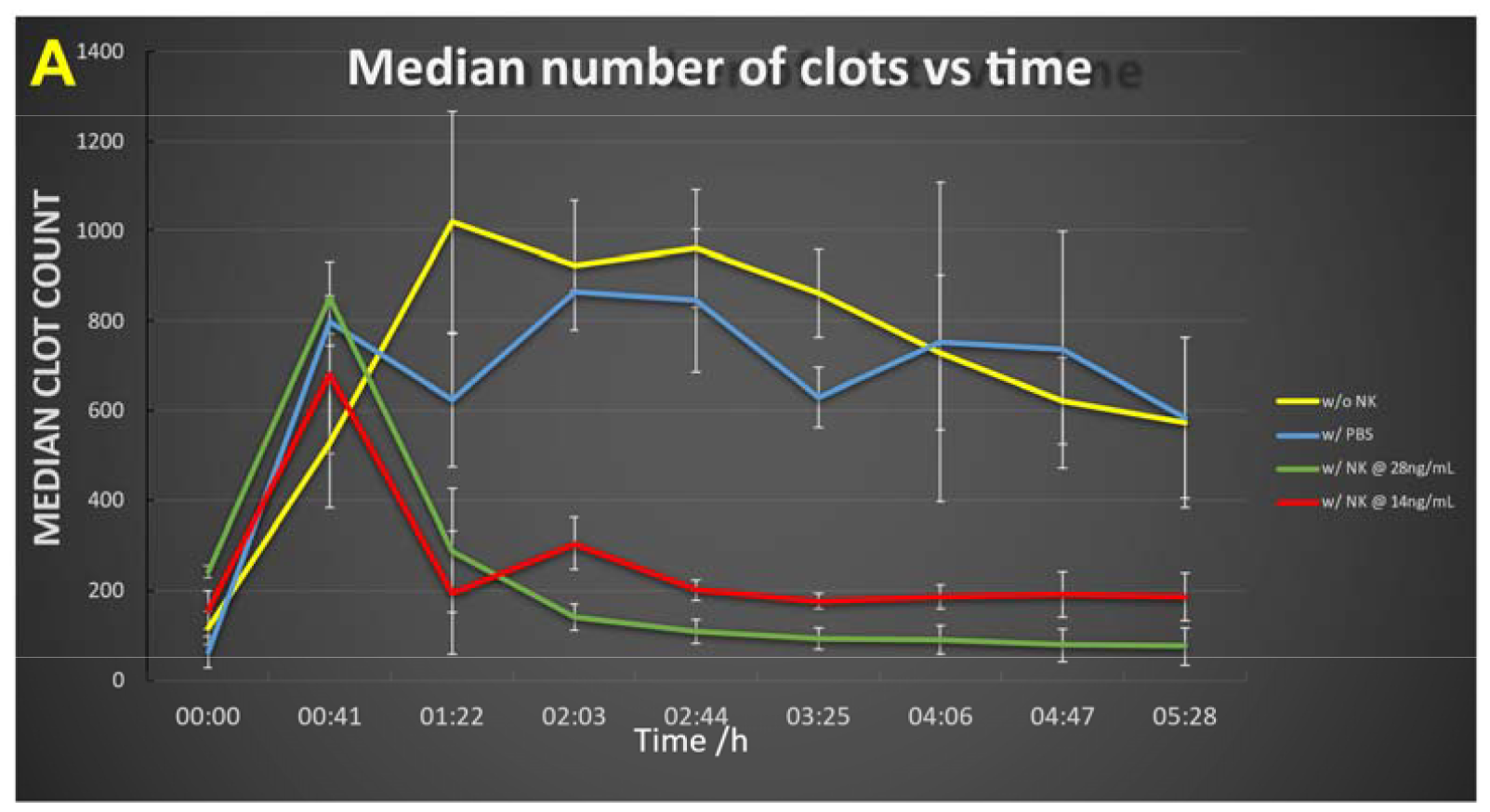
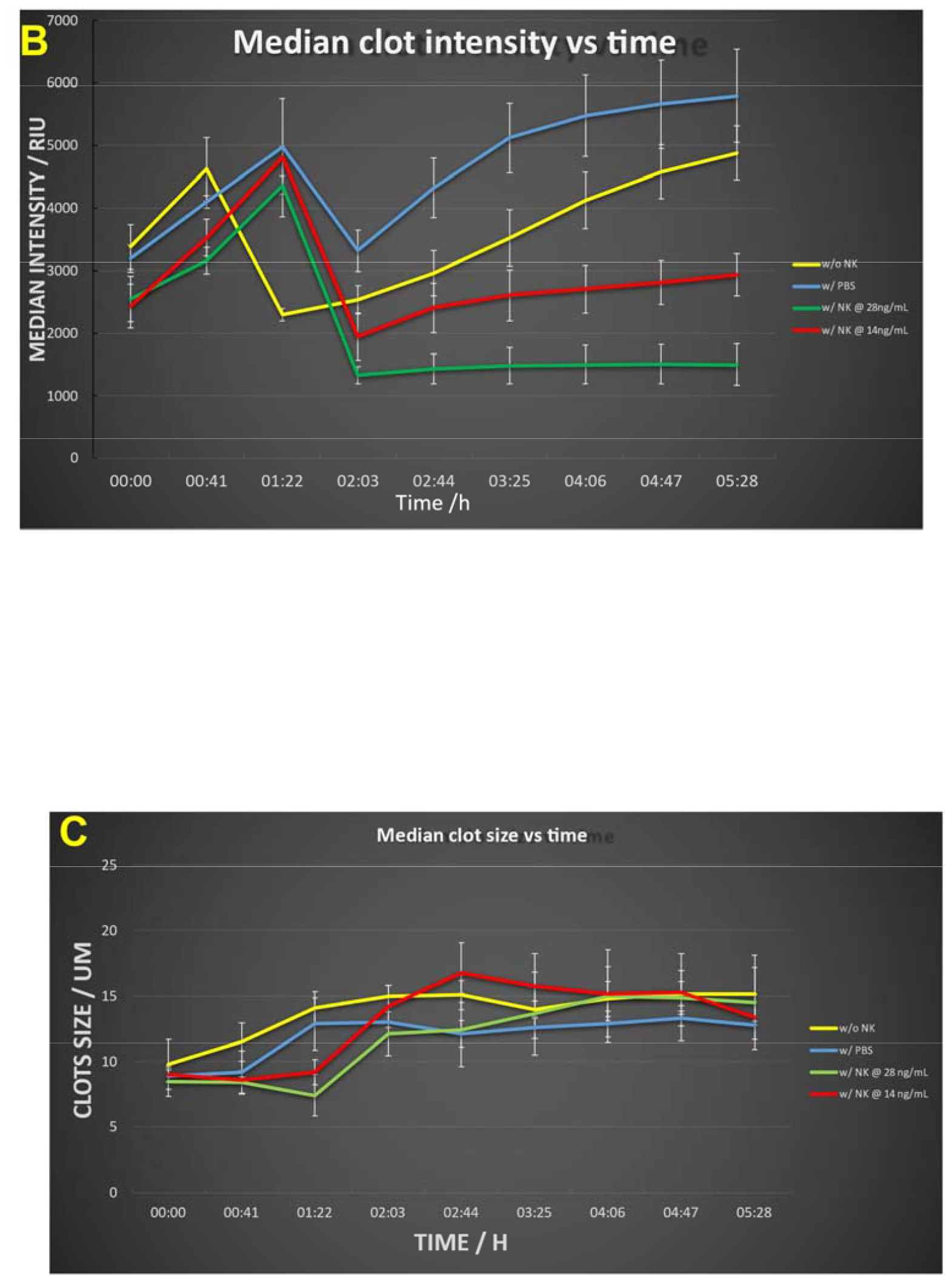
Time evolution of (A) clot number, (B) intensity and (C) median clot size during the development of fibrinaloid microclots and their incubation with nattokinase. Thrombin and fibrinogen were incubated together with thioflavin T, and imaged after 6h in a Cytation 1, as described in Methods. Further additions were none (yellow), PBS (blue), recombinant nattokinase 28 ng/mL (green), or recombinant nattokinase 14 ng/mL (red). Videos of the incubation with PBS and with nattokinase are given in Supplementary Information.

Three features stand out. First, especially in the absence of NK (yellow trace), the clots increase in number and size over time (Figure 2A, 2B), illustrating how microclots may aggregate to form macroclots, as part of the normal amyloidogenic process (e.g. [173,212,215,239,265-275]). This kind of aggregation may be highly significant in stroke and myocardial infarctions, where clots may be far larger than the simple sloughing off of atherosclerotic plaques might reasonably create. Secondly, the enzyme effectively decreases the rate and extent of microclot formation, in rough proportion to the amount of enzyme (compare e.g. red and green traces at 5h). The lowest intensity point was observed in the interval 2-4h, implying a die-off in activity or instability of the enzyme over time. This is good, in that untrammelled fibrinolytic activity may not be of the greatest therapeutic benefit. Lastly, the <u>median</u> clot size (Fig 2C) increases briefly then stabilizes. This reflects the fact that smaller clots will tend to be degraded preferentially as their surface area per unit mass is significantly greater than that of larger clots. (It is not commonly recognised, but if one imagines two solid spheres, of which one is twice the diameter of the other, the degradation of a given (i.e. the same) mass in the two spheres leads to a loss in mass of just 12.5% of the larger sphere when the smaller one is completely degraded, and a loss in radius of the larger sphere that is less than 5% of its starting value. Consequently, although possibly at first glance surprising, this is, given the traces in Figures 2A and 2B, in fact the result expected for Figure 2C.)

### Using Amytracker dyes instead of ThT

Because it is valuable to have other dyes should one wish to use multiple wavelengths (as in [103]), we also assessed the red oligothiophene-class Amytracker™ dyes (Ebba Biotech) (see e.g. [101,103,105,276-283]). However, these gave highly anomalous traces in this system, and we suspect may have inhibited the nattokinase, so were not further pursued.

## 3. Discussion

The ability to assess the rate of fibrin amyloid formation and degradation noninvasively is highly desirable, as it precisely permits studies of the present type that can then be automated. While still not a high-throughput approach in the usual sense, this does provide a substantial advance in scoring fibrinaloid microclot formation that is both fully quantitative and without undue operator fatigue. This has allowed us, for the first time, to conclude at least three important features: (i) the formation kinetics of fibrin amyloid microclots in whole samples may be imaged noninvasively in an automated manner, (ii) such microclots can aggregate over time, and (iii) the fibrinaloid microclots may be degraded by nattokinase. This latter has significant therapeutic implications for those suffering from Long COVID and related disorders, as NK preparations are widely available commercially. Our approach also thus allows for the comparison of different preparations of NK. Future work could usefully include recombinant serrapeptase (NK/SP), lumbrokinase (NK/LK) and/or sequence variants of NK/SP made using the methods of synthetic biology [284], since both serrapeptase and lumbrokinase also have fibrinolytic and amyloid-degrading properties [60,285-296].

## 4. Materials and Methods

### Assay method

*In vitro* microclots were made by mixing commercially obtained fibrinogen (Sigma catalogue number 9001-32-5, at a final concentration of 2 mg/mL) and thrombin (Sigma, final amount 14U) with bacterial LPS (Sigma product code L2630-10MG) and used at a final concentration of 1ng / mL and incubated at 37ºC for 30 min. Samples were then exposed to the fluorogenic amyloid dye, Thioflavin T (ThT) (final concentration: 0.03 mM) for 20 mins (protected from light) at room temperature. Following incubation, 10 μL of the recombinantly produced nattokinase at different concentrations / PBS (control) were added. This was then followed by immediately pipetting 15 μL of assay sample onto a 15-well slide ‘angiogenesis’ glass bottom plate used without a lid (bidi: https://ibidi.com/chambered-coverslips/245--slide-15-well-3d-glass-bottom.html), reproduced in Figure 3, and without shaking (cf. [142,297-309]). The excitation wavelength band for ThT was set at 450 nm to 499 nm and the emission at 499 nm to 529 nm. Samples were viewed using Gen5 software on an Agilent BioTek Cytation 1 Cell Imaging Multimode Reader, essentially following the protocol developed and described by Dalton and colleagues [179]. The Cytation instrument is an automated fluorescence microscope with 8-bit intensity resolution in which an entire, large field of view can be constructed at high magnification by taking serial images and moving the stage automatically. With the 4x objective used, each final image (as in Figure 1) was composed of 1296 individual images. The typical file size of a final, stitched .tif image was 19 Mb. Each experiment was run multiple times, each time being in triplicate (three separate wells). Other relevant settings that we optimized for this assay were as follows: the Cytation 1 temperature was set at 37C, and images were taken every 41 minutes for 6 hours. The colour channel used was GFP 469,525 A fixed focal height setting, with a bottom elevation of 549 μm and 0 μm offset was selected. A Z-Stack montage of the entire well was applied, with a step size of 86.9 μm, and 12 slices. Samples were analysed using the Gen5 Image Prime 3.13.15 software supplied with the instrument, and the thresholds for minimum and maximum object (clots) size that could be detected were set at 5 and 500 μm, respectively.

**Figure 3.**
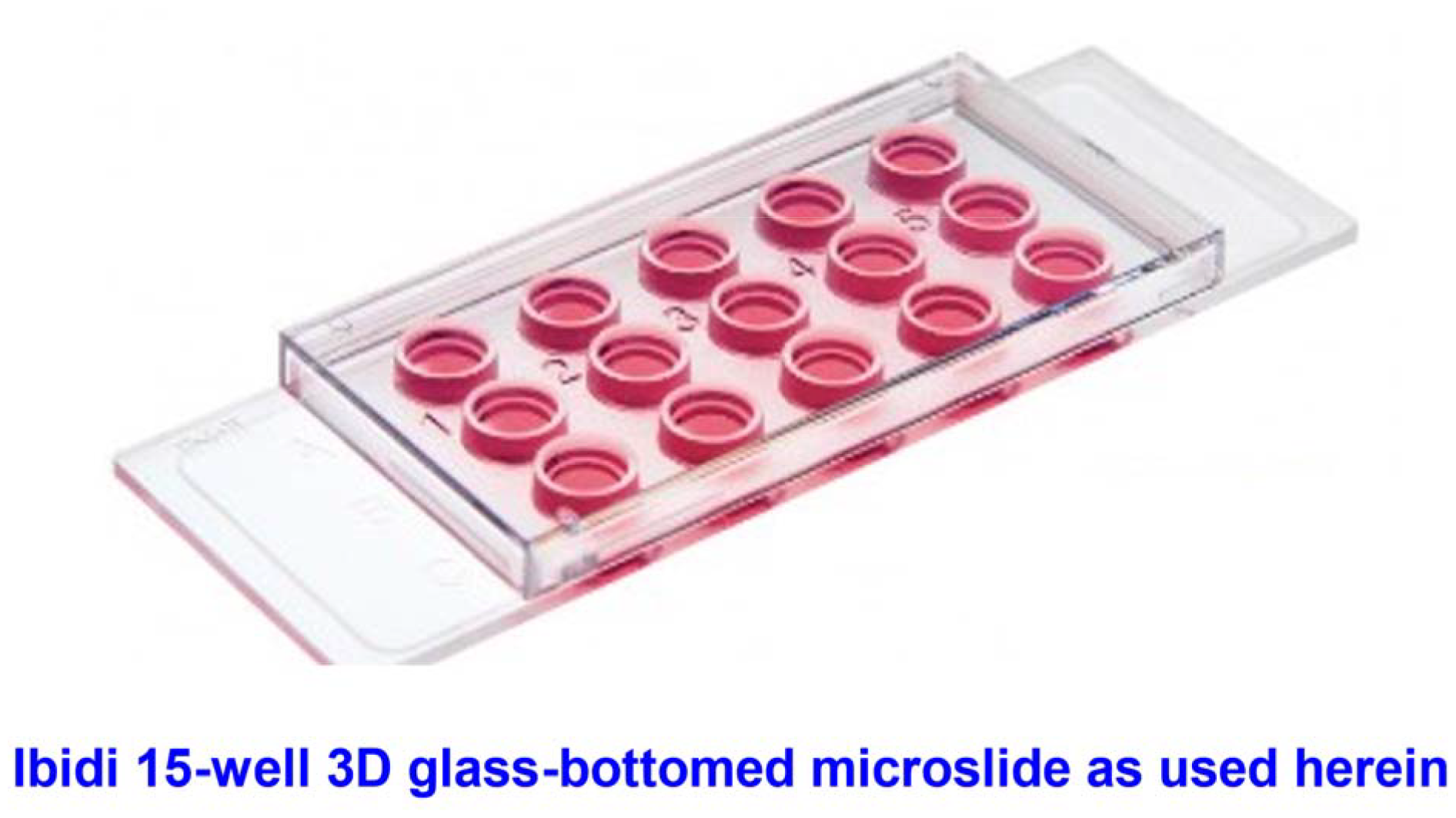
The incubation system used herein, allowing imaging from below

### Recombinant nattokinase

Recombinant nattokinase was produced within the Liverpool Gene Mill. The nucleotide sequence for *Bacillus subtilis* nattokinase (Uniprot <u>Q93L66</u>, GenBank: AER52006.1) was synthesised by Twist Bioscience and supplied in the pET28a(+) plasmid. The sequence was modified to include a C-terminal poly-Histidine tag for purification, as well as an N-terminal PelB leader sequence in which the terminal QPAMA residues are replaced by APOIA, and with a penta-aspartate linker for targeting to the periplasmic space [310] plus a ENLYFQ TEV cleavage site and a further SGS linker prior to the nattokinase sequence (beginning AQSVPY). The vector was used to transform chemically competent cells of the Rosetta™ strain of *Escherichia coli* (Novagen) according to the method described by Inoue et al [311]. Transformed cells were plated on plates of LB-agar (0.5% w/v yeast extract, 1% w/v NaCl, 1% w/v tryptone and 2% agar) supplemented with 50 µg/ml kanamycin and 25 µg/ml chloramphenicol. A single colony from the agar plate was used to inoculate 5 ml of LB broth (0.5% w/v yeast extract, 1% w/v NaCl, 1% w/v tryptone) supplemented with kanamycin and chloramphenicol as described above, for overnight culturing at 37°C with shaking. The culture was diluted to an OD_600_ of 0.05 in 500 ml of LB broth supplemented with kanamycin and chloramphenicol as described above, and incubated with shaking at 37°C. When an OD_600_ of 0.6 was reached, recombinant protein expression was induced by addition of 0.75 mM isopropyl β-D-1-thiogalactopyranoside (IPTG), and the culture was incubated overnight at 18°C with shaking. Cell pellets were harvested by centrifugation at 4000 xg for 10 minutes, and the pellets were resuspended in 50 ml of a solution of Tris-HCl (pH8) and 10 mM EDTA, and incubated at 60°C for 2 hours [312]. The suspension was centrifuged at 4°C at 16000 xg for 10 minutes, and the supernatant was passed through 1 ml of HisPur™ Ni-NTA resin (Thermo Scientific) to purify poly-Histidine-tagged proteins. Bound proteins were eluted using 500 mM imidazole, followed by desalting and concentration using a Pierce™ Protein Concentrator PES (Thermo Scientific) with 30 kDa cut-off. Protein yield was quantified using the Pierce™ Bradford Protein Assay Kit (Thermo Scientific), and samples were frozen with 10% v/v glycerol until further use. Inclusion body formation [313] was not here a significant issue. Figure 4 shows a gel illustrating the final preparation.

**Figure 4:**
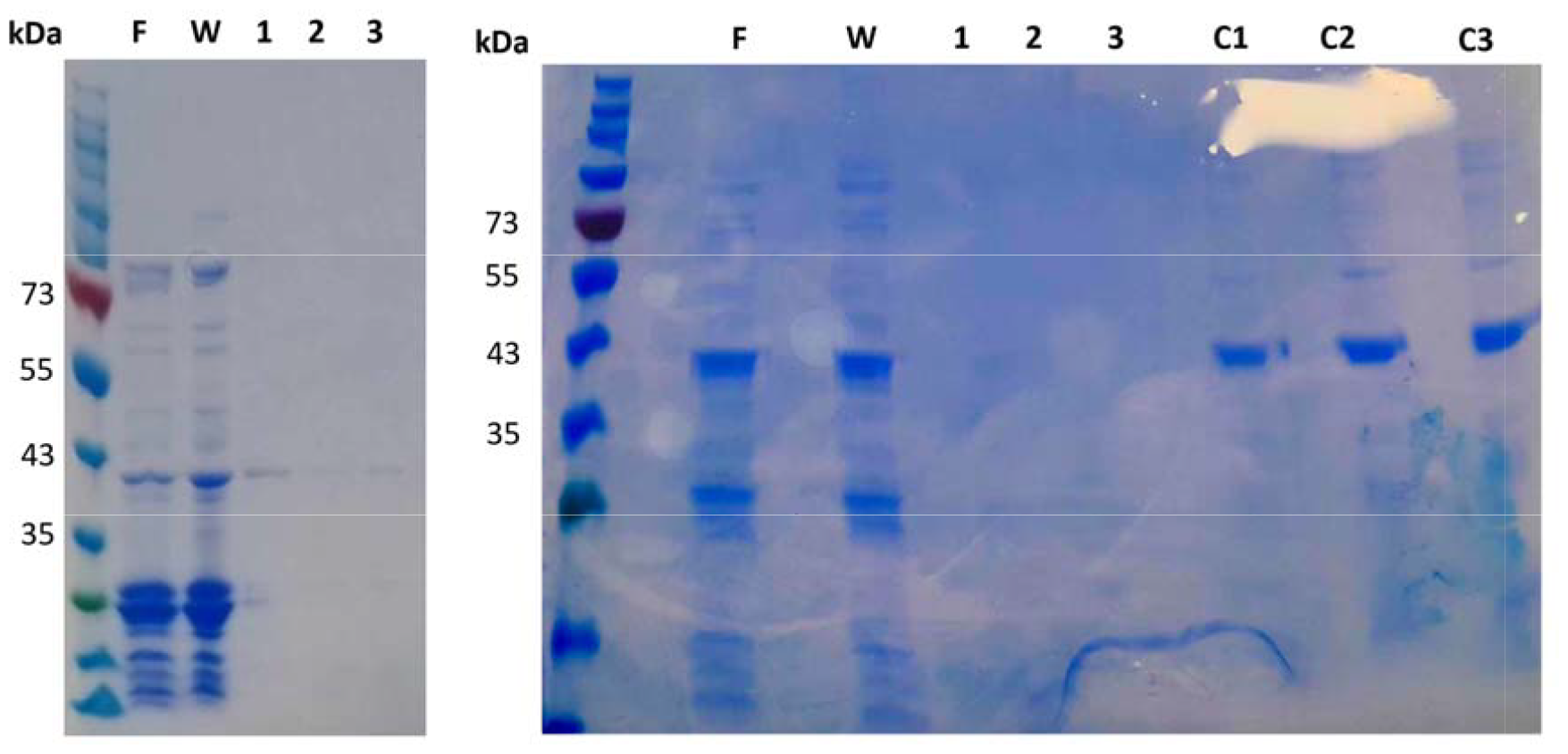
SDS-PAGE of recombinant Nattokinase. F - sample flow-through (unbound proteins), W — fraction of wash buffer (100 mM Tris, pH 7.5, 150 mM NaCI, 50 mM imidazole); 1-3 — purified fractions using elution buffer (100 mM Tris, pH 7.5,150 mM NaCI, 500 mM imidazole); C1&C2 - purified samples concentrated through 30 kD cut-off protein concentrator unit; C3 - C1 & C2 samples pooled and further concentrated through 3 kD cut-off protein concentrator unit.

A kinetic experiment was set up on the Cytation 1 and the effect of nattokinase on microclots was studied at final concentrations of 28 ng/μL and 14 ng/μL, using ThT at a final concentration of 0.005mM, as the fluorogenic dye. Measurements were taken every 40 minutes.

## Supplementary figures

Movies (not submitted in preprint) showing a kinetic series of images for samples treated with either PBS (Supplementary Figure 1) or 28 ng/mL nattokinase (Supplementary Figure 2). Frames start from read 1 at time zero and end at read 9 at time 5h 28m. Movies play at 0.5 frames per second. Time zero is the start of the reaction when PBS / NK was added to the sample containing fibrinogen, LPS, Thrombin, and ThT as described in the text. Green annuli are an artefact that may be ignored.

## Author Contributions

Conceptualization, DBK & EP; methodology, JMG, JES-S, CWT; resources, DBK & EP.; writing—original draft preparation, DBK; writing—review and editing, All authors; project administration, DBK & EP; funding acquisition DBK & EP. All authors have read and agreed to the published version of the manuscript.

## Acknowledgments

We thank Dr Caroline F. Dalton (Sheffield Hallam University) and Drs Amanda Barnes and Ashley Smith (Agilent) for many helpful discussions about the Cytation system and its optimisation

## Disclaimer/Publisher’s Note

The statements, opinions and data contained in all publications are solely those of the individual author(s) and contributor(s) and not of MDPI and/or the editor(s). MDPI and/or the editor(s) disclaim responsibility for any injury to people or property resulting from any ideas, methods, instructions or products referred to in the content.

